# Deficiency of DDI2 suppresses liver cancer progression by worsening cell survival conditions

**DOI:** 10.1101/2024.12.12.628086

**Authors:** Keli Liu, Shaofan Hu, Reziyamu Wufuer, Qun Zhang, Lu Qiu, Zhengwen Zhang, Meng Wang, Yiguo Zhang

## Abstract

The levels of reactive oxygen species (ROS) and the extent of ensuing DNA damage significantly influence cancer initiation and progression. Of crucial importance, the aspartate protease DDI2 has been proposed to play a pivotal role in monitoring intracellular ROS levels (to trigger oxidative eustress or distress), as well as in the oxidative DNA damage repair, through redox homeostasis-determining factor Nrf1 (encoded by *NFE2L1*). However, the specific role of DDI2 in the multi-step process resulting in the development and progression of liver cancer remains elusive to date. In the present study, we employed the CRISPR/Cas9 gene editing system to create two nuanced lines of *DDI2* knockout (i.e., *DDI2*^*−/−*^ and *DDI2*^*insG/−*^) from liver cancer cells. Subsequent experiments indicate that the knockout of *DDI2* leads to increased ROS levels in hepatoma cells by downregulating two major antioxidant transcription factors Nrf1 and Nrf2 (encoded by *NFE2L2*), exacerbating endogenous DNA damages caused by ROS and not-yet-identified factors, thereby inhibiting cell proliferation and promoting apoptosis, and ultimately hindering *in vivo* malignant growth of xenograft tumor cells. Conversely, the restoration of DDI2 expression reverses the accumulation of ROS and associated DNA damage caused by *DDI2* knockout, eliminating the subsequent inhibitory effects of *DDI2* deficiency on both *in vitro* and *in vivo* growth of liver cancer cells. Collectively, these findings demonstrate that *DDI2* deficiency impedes liver tumor growth by disrupting its survival environment, suggesting that DDI2 may serve as a novel therapeutic target for anti-cancer strategies aimed at modulating ROS or DNA damage processes.

## 1. Introduction

The liver is a vital organ responsible for metabolism, synthesis, detoxification, bile secretion and other functions within the body. However, it also serves, in the meantime, as the major site for the colonization of both primary and metastatic tumors ^1^. As a result, liver cancer is accepted as the leading cause of cancer-related deaths worldwide and thus ranks as the second most common cause in China (https://gco.iarc.fr/today/en). Of importance, the incidence of liver cancer continues to rise annually, but its prognosis remains poor ^2^. Therefore, further exploration of new therapeutic targets and improved strategies for treating liver cancer is essential for those extant and potential patients.

Clearly, oxidative stress generated by reactive oxygen species (ROS) ^3, 4^ and subsequent genomic instability resulting from oxidative DNA damage ^5, 6^ are viewed as significant factors influencing the initiation of cancer and ensuing progression of the malignant tumors, as bad as affecting the anti-cancer efficacy of relevantly targeted therapies. In liver cancer, ROS can impact tumor development and progression, and its chemoresistance by modulating cell processes of *e*.*g*., autophagy and metabolism ^7, 8^, whilst the DNA damage-related tumor progression is further determined primarily by those deficiencies in the DNA repair pathways and variations in the expression of DNA repair enzymes ^9^. Notably, ROS can indeed induce DNA damage and genomic instability, leading to the accumulation of carcinogenic alterations ^10^. In addition to the well-known transcription factor Nrf2 (NFE2-like bZIP transcription factor 2, encoded by *NFE2L2*), another family member Nrf1 (NFE2-like bZIP transcription factor 1, encoded by *NFE2L1*) also plays a crucial role in regulating robust redox homeostasis to monitor intracellular ROS levels, and hence influencing tumor development and progression ^11, 12^. Of note, the aspartyl protease DDI2 (i.e., DNA damage inducible 1 homolog 2) has been identified as a key cytosolic protease in the processing and activation of the endoplasmic reticulum (ER) membrane-bound Nrf1. This is because it enables for a proteolytic cleavage within the N-terminal domain of ubiquitinated and deglycosylated Nrf1 to become a soluble, activated isoform, hence facilitating its entry into the nucleus for its transcriptional regulation of proteasome subunits, oxidative stress-defending genes, and other cytoprotective target genes ^13, 14^. Thereby, the DDI2-Nrf1-target gene network plays a significant regulatory role in the sensitivity of tumors to proteasome inhibitors as used in relevant cancer treatment ^15, 16^. Furthermore, DDI2 also influences the process of DNA damage and affects genome stability by modulating the DNA replication stress response ^17, 18^. These prior studies suggest that DDI2 may exert a significant regulatory potential in tumor progression by regulating both ROS levels and ensuing DNA damage processes. However, DDI2 remains to be a poorly understood protease, so that its regulatory effects on oxidative stress, DNA damage, and tumor progression in liver cancer are not yet well elucidated.

In this study, we investigated the regulatory effects of DDI2 on the biological behavior of liver cancer cells, both *in vitro* and *in vivo*, by examining two distinct *DDI2* knockout (i.e., *DDI2*^*−/−*^ and *DDI2*^*insG/−*^) hepatoma cell lines. The experimental results demonstrated that knockout of *DDI2* leads to increased ROS levels and exacerbates DNA damage in hepatoma cells. This, in turn, inhibits the proliferation and migration of tumor cells, but promotes apoptosis, ultimately resulting in the suppression of xenograft tumor growth *in vivo*. Furthermore, restoring DDI2 expression significantly counteracts the inhibitory effects of *DDI2* knockout on the malignant growth of liver cancer in both *in vitro* and *in vivo* contexts. Collectively, these findings suggest that DDI2 is likely to serve as a potential target for targeted anti-cancer therapeutic strategies aimed at modulating redox homeostasis and DNA damage repair system.

## 2. Materials and Methods

### 2.1. Cell culture and treatments

HepG2 cells and HEK293T cells were originally derived from American Type Culture Collection (ATCC, Manassas, VA, USA). Mycoplasma testing was performed prior to experimentation. Both cell lines were saved in DMEM supplemented with 5 mM glutamine, 10% (v/v) fetal bovine serum (FBS), and 100 units/mL of either penicillin or streptomycin in a 37°C incubator with 5% CO_2_. Additionally, two distinct HepG2-derived cell lines with *DDI2* knockout, referred to as *DDI2*^*−/−*^ and *DDI2*^insG/*−*^ (with an additional G inserted to yield a putative unusual peptide by shifting its reading frame as shown in Figure S1), were herein established through CRISPR/Cas9-editing of *DDI2* using specific gRNA (in Table S1). The authenticity of these *DDI2* knockout cells was confirmed to be true through authentication analysis. Thereafter, experimental cells were transfected with a Lipofectamine 3000 mixture containing a DDI2-expressing construct or another control vector for 8 h, followed by an additional 24-h recovery period in fresh medium prior to experimentation. Furthermore, additional experimental cells were treated, respectively, with the DNA damage inducer cisplatin (referred to as DDP, 15663-27-1, Aladdin, Shanghai, China), the ROS inhibitor N-Acetylcysteine (referred to as NAC, A601127, Sangon Biotech, Shanghai, China), or the proteasomal inhibitor MG132 (M7449, Sigma Aldrich) for distinct lengths of time as indicated in relevant figure legends.

### 2.2. Expression constructs

An expression construct for human DDI2 was created by subcloning its full-length cDNA sequence into the pcDNA3 vector, utilizing a pair of forward and reverse primers (as listed in Table S1), which were synthesized by Sangon Biotech Co. (Shanghai, China). The fidelity of all constructs was further confirmed to be true through sequencing.

### 2.3. Western blotting (WB) with distinct antibodies

Total cell lysates in a lysis buffer (0.5% SDS, 0.04 mol/L DTT, pH 7.5), supplemented with protease and phosphatase inhibitors (each comprising cOmplete and PhosSTOP EASYpack tablets in 10 mL buffer), were denatured immediately at 100°C for 10 min, sonicated adequately, and then diluted in 3× loading buffer (187.5 mmol/L Tris-HCl, pH 6.8, 6% SDS, 30% Glycerol, 150 mmol/L DTT, 0.3% Bromphenol Blue) at 100°C for 5 min. Subsequently, equal amounts of protein extracts were subjected to separation by SDS-PAGE with a 4-15% gradient polyacrylamide gel, followed by visualization through immunoblotting with specific antibodies (as indicated in Table S1). Some of the blotted membranes were stripped for 30 min and re-probed with additional primary antibodies. Therein, β-actin served as an internal control to verify the equal loading of proteins.

### 2.4. Quantitative real-time PCR (qPCR)

Approximately 500 ng of total RNAs extracted from experimental cells was subjected to a reverse-transcription reaction to generate the first strand of cDNA. The newly synthesized cDNA served as the template for real-time qPCR in the Master Mix, which was subsequently deactivated at 95°C for 10 min. This was followed by 40 cycles of amplification, comprising an initial annealing at 95°C for 15 s, and an extension step at 60°C for 30 s. The final melting curve was analyzed to assess the quality of amplification. The β-actin mRNA level was utilized as an optimal internal standard control, and thus relevant epression levels of target genes were quantified via real-time qPCR, using each pair of the indicated primers (as listed in Table S1). The resulting data were shown graphically as fold changes (mean ± S.D.) relative to the control values, which were obtained from at least three different experiments performed each in triplicates.

### 2.5. Dual-Luciferase reporter gene assay

After allowing HEK293T cells (1.0 × 10^5^) to grow in each well of 12-well plates until they reached 80% confluence, the cells were co-transfected with a Lipofectamine 3000 mixture containing either *pNrf1-luc* or *pNrf2-luc* established by Qiu *et al* ^19^, along with a DDI2 expression plasmid. In this dual reporter assay, the *Renilla* expression driven by pRL-TK served as an internal control for transfection efficiency. The resulting data were normalized based on at least three independent experiments, each of which was performed in triplicate, and are presented as a fold change (mean ± S.D.) relative to the control values.

### 2.6. Detection of ROS, cell cycle and apoptosis by flow cytometry

Equal numbers of experimental cells (3 × 10^5^ cells in each well of 6-well plates) were allowed to grow for 24 h after transfection with a DDI2 expression or empty control plasmid. Once the cells reached 80% confluence, intracellular ROS levels were measured according to the instruction as provided in the ROS assay kit (S0033S, Beyotime, Shanghai, China). Additionally, cell cycle and apoptosis were also determined following the protocols of the relevant kits (C1052 and C1062S, Beyotime, Shanghai, China). Subsequent analyses were performed using the CytoFLEX instrument (Beckman Coulter, Ltd., Miami, FL, USA) and FlowJo software (version 10.3.0, Tree Star, Ltd., USA). The data are graphically shown as fold changes (mean ± S.D.) relative to their control values.

### 2.7. Transcriptome sequencing and bioinformatics Analysis

Equal numbers of wild-type (*WT*) HepG2, its derived *DDI2* knockout cell lines *DDI2*^*−/−*^ and *DDI2*^insG/*−*^ were seeded in each well of 6-well plates. Once cell confluence reached 80%, the cells were harvested and total RNAs were extracted using 500 μL TRNzol Universal (#DP424, TIANGEN, Beijing, China). The quality control was performed using the Agilent2100 with criteria of RNA ≥ 200 ng, 28S/18S ≥ 1.0 and RIN ≥ 7.0. Then, cDNA libraries were constructed for its RNA-seq using the DNBSEQ platform (BGI, China; contract No. F21FTSCCKF4365_PEOoawhR). The clean reads were filtered and aligned to the reference genome (GRCh38). The mRNA expression levels of interested genes were normalized using the FPKM method, and those differentially expressed mRNAs were identified by using the Dr.Tom online analysis platform (BGI, China). The threshold for differentially expressed genes (DEGs) were defined by their “Q-value (adjusted p-value) ≤ 0.05 and the absolute value of |log2 (Fold Change)| ≥ 1”. Distinct expression heatmaps for those selected DEGs were generated using the pheatmap package in R (version 4.3.0), in which divergent gene expression trends across distinct samples were evinced in different colors, with red indicating higher expression levels and green indicating lower expression levels. Additionally, the Gene Set Enrichment Analysis (GSEA) analysis of cell cycle pathway was also conducted using GESA software (version 4.0.3).

### 2.8. Immunocytochemistry of γ-H2AX by confocal microscopy

Each group of experimental cells (3 × 10^5^) was allowed to grow in individual wells of 6-well plates. After the cells fully adhered, they were treated with or without 10 μM DDP for 22 h and subsequently washed with PBS before being assessed for DNA damage. The fixed cells were blocked and then incubated with an antibody against γ-H2AX (C2035S, Beyotime, Shanghai, China), followed by their DNA staining with DAPI (4’,6-diamidino-2-phenylindole), according to the manufacturer’s instructions (I029-1-1, Nanjing Jiancheng, Nanjing, China). The immunofluorescently stained cells were subjected to confocal microscopic observation, by detecting green fluorescent γ-H2AX at λ ex/em = 495/519 nm and blue fluorescent DAPI at λ ex/em = 464/454 nm.

### 2.9. Detection of 8-Hydroxy-2-deoxyguanosine (8-OHdG)

The yield of 8-Hydroxydeoxyguanosine (8-OHdG, as a well-known marker of oxidative stress) in *WT, DDI2*^*−/−*^ and *DDI2*^insG/*−*^ cell lines was measured by using an 8-OHdG-relevant ELISA Kit (E-EL-0028, Elabscience, Wuhan, China). Briefly, genomic DNAs were extracted from experimental cells using the Qiagen’s Genomic-tips 20/G, the Genomic DNA buffer set and proteinase k, and then stored at −20°C for later quantification of 8-OHdG. To measure 8-OHdG, DNAs were bound to a 96-well flat-bottom plate, followed by the first wash before addition of the capture antibody. After a second wash, the detection antibody and an enhancer solution were added. The color-developing solution was also subsequently added, and the resulting absorbance was measured at 450 nm by using the Biotek Synergy HTX Multi-mode microplate reader.

### 2.10. The CCK8 Assay for cell proliferation

The rate of cell proliferation was measured using the cell counting kit 8 (CCK8, Biosharp, China). Briefly, the experimental cells were seeded at a density of 4000 cells per well in 96-well plates. Subsequently, 10 μL of the CCK8 reagent was added to each well of the 96-well plates before being incubated for 2 h at 37°C in a 5% CO_2_ atmosphere. Lastly, the optical densities of the wells were measured using a microplate reader at 450 nm. It is important to note that the entire experimental process was conducted in the absence of light.

### 2.11. Transwell-based migration assay

When the growing cells reached 70% confluency, they were subjected to a 12-h starvation period in serum-free medium. The experimental cells (5 × 10^3^) were suspended in 0.5 mL of medium containing 5% FBS and then seeded in the upper chamber of a transwell. This setup allows the cells to grow on a microporous polycarbonate membrane that has been tissue culture-treated to enhance cellular attachment to the bottom. The cell-seeded transwells were subsequently placed in each well of 24-well plates containing 1 mL of complete medium (i.e., the lower chamber), and cultured for 24 h in the incubator at 37°C with 5% CO_2_. Thereafter, the remaining cells in the upper chamber were removed, before those cells, that had attached to the lower surface of the transwell membranes, were fixed with 4% paraformaldehyde (AR10669, BOSTER) and stained with 1% crystal violet reagent (Sigma) prior to being counted.

### *2*.*12. In vitro* scratch assay

Experimental cells (1 × 10^5^) were allowed for growth in 6-well plates to reach 70% confluency, before being synchronized by 12-h starvation in serum-free medium and additional 6-h treatment with 1 μg/mL of mitomycin C (Cayman, USA). Thereafter, a clear ‘scratch’ was created in the cell monolayer, and then allowed to heal in continuously culturing at 37°C with 5% CO_2_. The wound closure distance was measured at the beginning (*T*_*0*_) and end of the experiment (*T*_*X*_). The formula (*T*_*0*_ *– T*_*X*_)/ *T*_*0*_ × 100 was used to calculate a percentage (%) of the wound closure by the migrated area for the ‘scratch’ to be healed.

### 2.13. Subcutaneous tumor xenograft model

Mouse xenograft tumor models were established by subcutaneously heterotransplanting human HepG2 *WT, DDI2*^*−/−*^ or *DDI2*^insG/*−*^ cell lines. Briefly, equal amounts (1 × 10^7^) of cells in the exponential growth phase were suspended in 0.1 ml of serum-free medium and inoculated subcutaneously at a single site in the right upper back region of male nude mice (BALB/C nu/nu, 6 weeks, 18-20 g). The injection procedure of all mice was completed within 30 min. Subsequently, the formation of murine subcutaneous tumor xenografts was monitored and their sizes were measured until the mice were sacrificed. These transplanted tumors were excised immediately after euthanasia and their sizes were calculated using a standard formulate (V = ab^2^/2). All mice were maintained under standard animal housing conditions with a 12-hour dark cycle and provided ad libitum access to sterilized water and diet, in accordance with the institutional guidelines for the care and use of laboratory animals (license SCXK-PLA-20210211). All these animal experimental procedures had been approved from the Ethics Committee of Chongqing Medical University.

### 2.14. H&E staining and immunohistochemistry

The xenograft tumor tissues were immersed in 4% paraformaldehyde overnight before being transferred to 70% ethanol. Individual tumor tissues were placed in processing cassettes, dehydrated through a series of alcohol gradient, and embedded in paraffin wax blocks. The paraffin-embedded samples were then sectioned to yield a series of 5-μm-thick slides. The tissue sections were de-waxed in xylene, rehydrated through decreasing concentrations of ethanol, and washed in PBS. Subsequently, they were stained by routine hematoxylin and eosin (H&E) and visualized using microscopy. For immunohistochemical analysis, the slides of tumor tissues were de-paraffinized in a xylene solution and subsequently dehydrated in concentration-graded ethanol before the inactivation of endogenous peroxidase activity. The samples were then subjected to being, in a microwave, boiled for 15 min in a citrate buffer (pH 6.0) to retrieve antigens, followed by being blocked with 1% BSA for 60 min. Thereafter, the sample sections were incubated with primary antibodies against caspase-9 (CASP9) or cyclin D1 (CCND1) at 4°C overnight, and then re-incubated for 60 min with a biotin-conjugated secondary antibody at room temperature before being visualized by DAB staining. The resultant images were acquired under a light microscope (Leica DMIRB, Leica, Frankfurt, Germany) equipped with a DC350F digital camera.

### 2.15. Statistical analysis

Significant differences were statistically determined using the Student’s *t-*test and one-way analysis of variance (ANOVA), except for somewhere indicated. The data are herein shown as a fold change (mean ± S.D., relative to corresponding controls), each of which represents at least three independent experiments that were each performed in triplicate.

## 3. Results

### 3.1. Deficiency of *DDI2* causes accumulation of ROS by downregulating antioxidant-related genes

To gain an insight into the role of DDI2 in liver cancer, herein we employed CRISPR/Cas9 gene editing technology to create a monoclonal knockout cell line in which *DDI2* is genomically effectively deleted in HepG2 cells. As a consequence, two distinct monoclonal cell lines were identified by genome sequencing of *DDI2* and verification of its expression (Fig. 1, A & B), and respectively designated as *DDI2*^*−/−*^ (with successful knockout of this gene-specific locus) and *DDI2*^insG/*−*^ (with an extra G inserted to shift its reading frame and yield a putative unusual peptide) (Fig. S1, A & B). Given that DDI2 is a crucial enzyme involved in the activation of the antioxidant regulatory factor Nrf1, hence it is initially hypothesized that the knockout of *DDI2* could induce oxidative stress in relevant tumor cells. Through Western blotting and real-time qPCR experiments, we assessed the expression of antioxidant-related genes and also observed a significant reduction in the expression of the antioxidant transcription factors Nrf1 and Nrf2, as well as their downstream antioxidant genes, due to knockout of *DDI2* (Fig. 1A, B). Furthermore, such knockout of *DDI2* also led to significant decreases in the expression levels of the core proteasomal subunits (PSMB5, PSMB 6 and PSMB 7) in the response to reduction of both Nrf1 and Nrf2 (Figs. 1, A & B, and S2A). Conversely, overexpression of DDI2 enabled to enhance the transcriptional activity of each of Nrf1 and Nrf2 promoters-driven luciferases (Fig. S2, B & C). Since previous researches conducted by our group and others had highlighted the significant regulatory roles of Nrf1 and Nrf2 in the progression of liver cancer ^19, 20^, in this study we further analyzed the correlation between the expression of DDI2 and these two major CNC-bZIP factors in clinical liver cancer samples. By utilizing the TCGA-LIHC data from the UCSC Xena database ^21^, the results revealed that the expression of DDI2 is significantly positively correlated with the expression of both Nrf1 and Nrf2 (Fig. 1C). Such positive correlation of DDI2 with Nrf1 and Nrf2 has been also further corroborated by bioinformatics using other distinct databases e.g., GEPIA2 ^22^, TIMER ^23^, and UALCAN ^24^ (Fig. S2D). After confirming downregulation of the antioxidant transcription factors Nrf1 and Nrf2 by knockout of *DDI2*, we examined the changes in ROS levels in liver cancer cells following *DDI2* knockout by employing flow cytometry and immunofluorescence experiments. As anticipated, these experimental results unraveled significant increases in intracellular ROS levels after knockout of *DDI2* in tumor cells (Fig. 1, D & E). Furthermore, we further performed transcriptomic sequencing on such *DDI2* knockout cells. As a result, an expression heatmap for antioxidant-related genes was generated by bioinformatics statistical analysis of the global profile and differentially expressed genes, which demonstrated an overall downward trend in these genes regulated by knockout of *DDI2* (Fig. 1F and S3).

**Figure 1.**
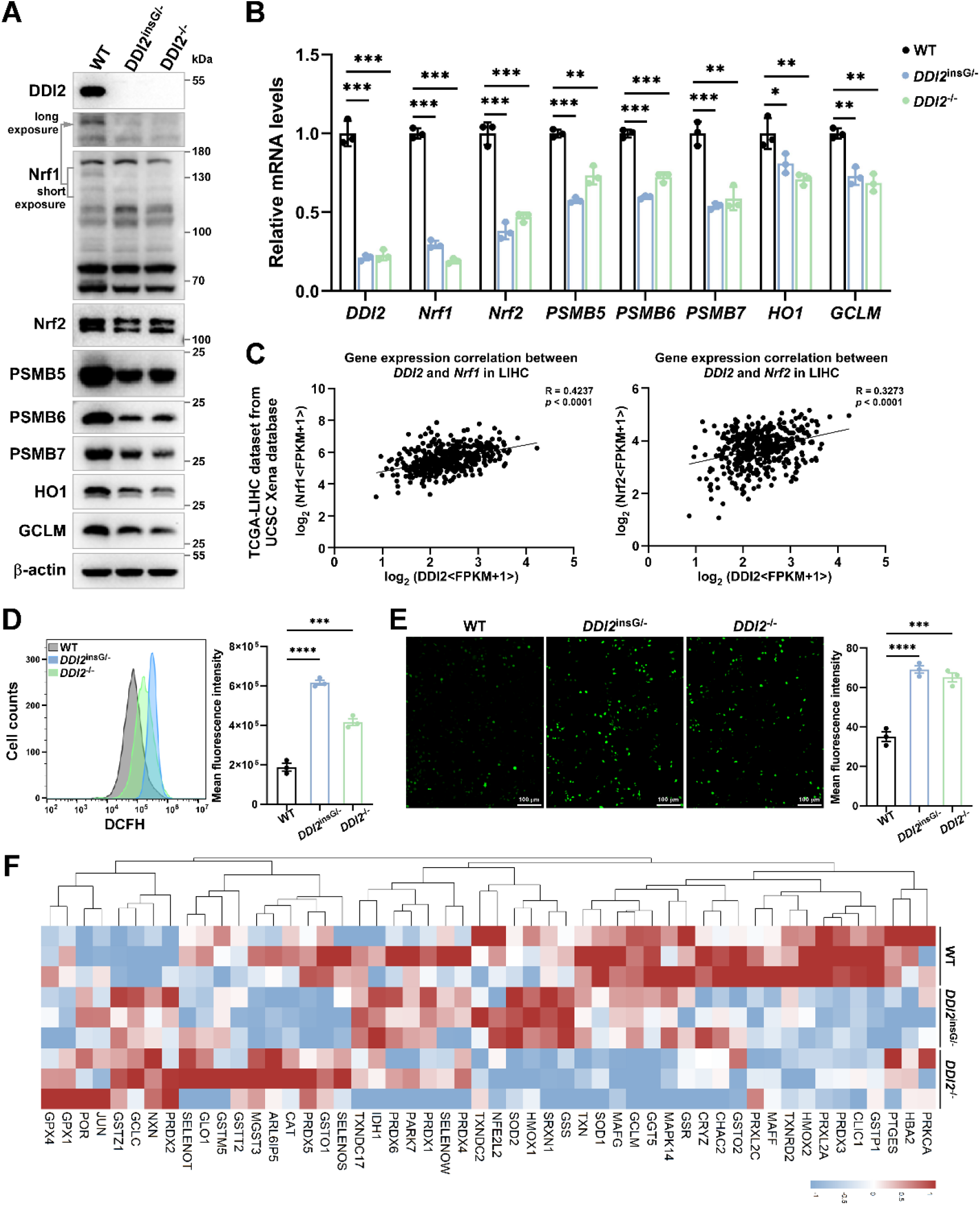
Knocking out *DDI2* leads to ROS accumulation by downregulating antioxidant-related genes. (A) Wild-type HepG2 cells (WT) and HepG2-derived *DDI2* knockout cells (designated as *DDI2*^insG/-^ and *DDI2*^-/-^) were subjected to Western blotting using the indicated antibodies. (B) Wild-type HepG2 cells (WT) and HepG2-derived *DDI2* knockout cells underwent real-time qPCR analysis to assess the mRNA expression levels of specified antioxidant-related genes. The results are presented as fold changes (mean ± S.D., n = 3), with significant differences indicated (*, p < 0.05; **, p < 0.01; ***, p < 0.001) relative to the WT controls. (C) The correlation of gene expression between DDI2 and Nrf1 or Nrf2 was analyzed using the TCGA-LIHC dataset from the UCSC Xena database. (D, E) Flow cytometry (D) and immunofluorescence (E) were employed to measure ROS levels in wild-type HepG2 cells (WT) and *DDI2* knockout cells (*DDI2*^insG/-^ and *DDI2*^-/-^), with the mean fluorescence intensity of ROS calculated. The results are expressed as mean fluorescence intensity (mean ± S.D., n = 3), with significant changes denoted (***, p < 0.001; ****, p < 0.0001) relative to the WT controls. (F) A differential expression heatmap of antioxidant-related genes in wild-type HepG2 (WT) cells and *DDI2* knockout cells (*DDI2*^insG/-^ and *DDI2*^-/-^) was generated based on transcriptome sequencing analysis.

### 3.2. Knockout of *DDI2* promotes DNA damage in liver cancer cells

Since the above evidence revealed that ROS accumulation results from knockout of *DDI2*, we thus further examined the regulatory effect of DDI2 on the DNA damage response in liver cancer cells by fluorescence detection and protein expression analysis of γ-H2AX (a phosphorylated form of histone H2A.X that exerts a crucial role in the DNA damage response). This is just because increased levels of γ-H2AX results from DNA damage, leading to enhanced green fluorescence of this marker primarily in the nucleus of *DDI2*-deficient cell lines (Fig. 2A); such changes in its fluorescence intensity serve as an intuitive indicator of the extents of DNA damage). As anticipated, our experimental results revealed that knockout of *DDI2* significantly promotes DNA damage in liver cancer cells (Fig. 2B). As a well-known inducer of DNA damage, Cisplatin (also referred to as DDP) was administrated in the relevant experiments. The results, as depicted in Figure 2 (A to C), indicate that deficiency of *DDI2 de facto* exacerbates the extents of DNA damage induced by cisplatin in *DDI2*^insG/*−*^ and *DDI2*^*−/−*^ cell lines when compared to *WT* cells.

**Figure 2.**
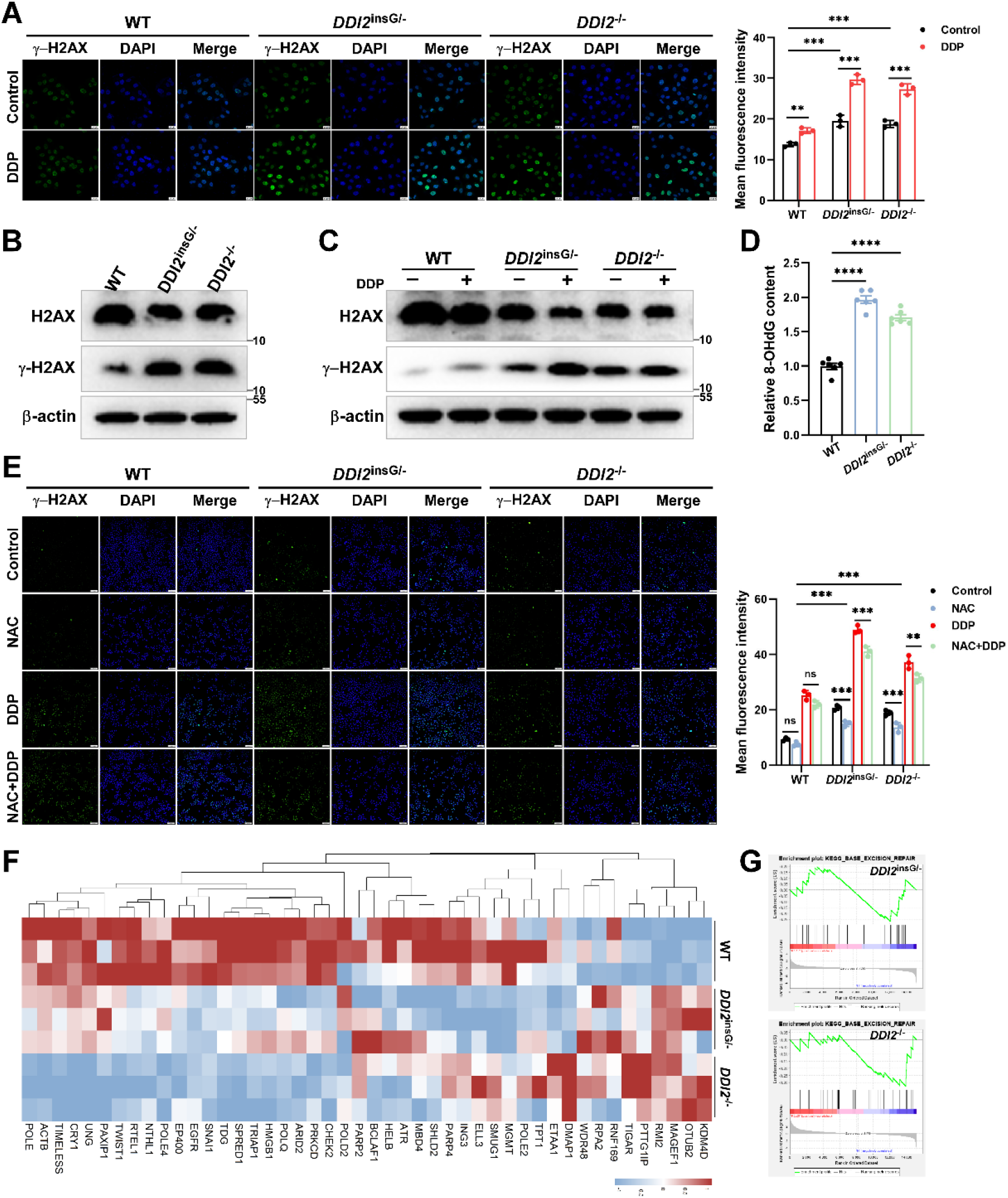
*DDI2* knockout exacerbates oxidative and other types of DNA damage. (A) Wild-type HepG2 cells (WT) and HepG2-derived *DDI2* knockout cells (designated as *DDI2*^insG/-^ and *DDI2*^-/-^) were treated with 10 μM DDP for 22 h and subsequently analyzed for the fluorescence intensity of γ-H2AX via immunofluorescence. The mean fluorescence intensity of γ-H2AX was calculated, with results expressed as mean fluorescence intensity (mean ± S.D., n = 3), and significant changes indicated (**, p < 0.01; ***, p < 0.001) relative to the controls. (B) Wild-type HepG2 cells (WT) and *DDI2* knockout cells (*DDI2*^insG/-^ and *DDI2*^-/-^) were subjected to Western blotting to assess the protein levels of H2AX and γ-H2AX. (C) Wild-type HepG2 cells (WT) and *DDI2* knockout cells (*DDI2*^insG/-^ and *DDI2*^-/-^) underwent treatment with 10 μM DDP for 22 h, followed by Western blotting to evaluate the abundance of H2AX and γ-H2AX. (D) The content of 8-OHdG was assessed in wild-type HepG2 cells (WT) and *DDI2* knockout cells (*DDI2*^insG/-^ and *DDI2*^-/-^) using the 8-OHdG DNA Damage Quantification Direct Kit. Results are presented as fold changes (mean ± S.D., n = 6), with significant differences indicated (****, p < 0.0001) relative to the WT controls. (E) Wild-type HepG2 cells (WT) and *DDI2* knockout cells (*DDI2*^insG/-^ and *DDI2*^-/-^) were treated with 10 μM DDP for 22 h, with or without the addition of 5 mM NAC, which was administered one hour prior to DDP treatment and maintained for a total 23 h. Following these treatments, the fluorescence intensity of γ-H2AX was assessed using immunofluorescence. The mean fluorescence intensity of γ-H2AX was calculated and expressed as mean fluorescence intensity (mean ± S.D., n = 3), with significant changes denoted (ns, non-significant; **, p < 0.01; ***, p < 0.001) relative to the controls. (F) A differential expression heatmap of DNA damage repair-related genes in wild-type HepG2 cells (WT) and *DDI2* knockout cells (*DDI2*^insG/-^ and *DDI2*^-/-^) was generated based on transcriptome sequencing analysis. (G) Gene Set Enrichment Analysis (GSEA) of the base excision repair pathway in *DDI2*^insG/-^ (NES = -1.129, p = 0.035) and *DDI2*^-/-^ (NES = -1.311, p = 0.0013) cells was conducted in comparison to WT cells, utilizing transcriptome sequencing data.

Next, we examined changes in the content of 8-hydroxy-2’-deoxyguanosine (8-OHdG), which acts as a key biomarker of oxidative damage to DNA due to reactive oxygen and nitrogen species (ROS/RNS), and hence reflects the level of DNA oxidative damage. The results revealed that knockout of *DDI2* further promotes oxidative DNA damage in *DDI2*^insG/*−*^ and *DDI2*^*−/−*^ cell lines, when compared to *WT* controls (Fig. 2D). Subsequently, we further assessed changes ofγ-H2AX in the DNA damage extents after inhibition of oxidative stress by the ROS inhibitor N-Acetylcysteine (NAC). As expected, the results demonstrated that whilst NAC can mitigate oxidative DNA damage to certain extents, such endogenous DNA damage arising by knockout of *DDI2* remains to be mostly exacerbated to rather higher extents in *DDI2*^insG/*−*^ and *DDI2*^*−/−*^ cell lines, particularly after being co-treated with DDP (Fig. 2E). From this, it is inferable that such *DDI2*-deficient DNA damage encompasses not only ROS-related type of damage and also other forms of DNA damage resulting from knockout of *DDI2*. Thereby, an expression heatmap analysis along with another gene set enrichment analysis (GSEA) on DNA damage repair-related regulatory genes was conducted according to the transcriptomic sequencing data, coincidentally revealing a downward trend in most of those DNA repair gene expression in such *DDI2*-deficient cell lines compared to *WT* cells (Fig. 2, F & G).

### 3.3. Knockout of *DDI2* diminishes both proliferation and migration but promotes apoptosis of liver cancer cells *in vitro*

Given that *DDI2* knockout promotes oxidative stress and DNA damage in hepatoma cells, it is much likely to impact on the cell survival. Thereby, we herein investigated whether *DDI2* knockout has an effect on the *in vitro* behavior of liver cancer cells. By flow cytometry and CCK8 assay to assess cell cycle progression and proliferation activity, respectively, it was found that *DDI2* knockout led to a remarkable arrest of examined cell cycle at the G0/G1 phase but its S/G2 phases were significantly shortened by *DDI2*^insG/*−*^ or *DDI2*^*−/−*^ (Fig. 3A), such that the cell proliferation was inhibited by *DDI2* knockout, as a consequence (Fig. 3B).

**Figure 3.**
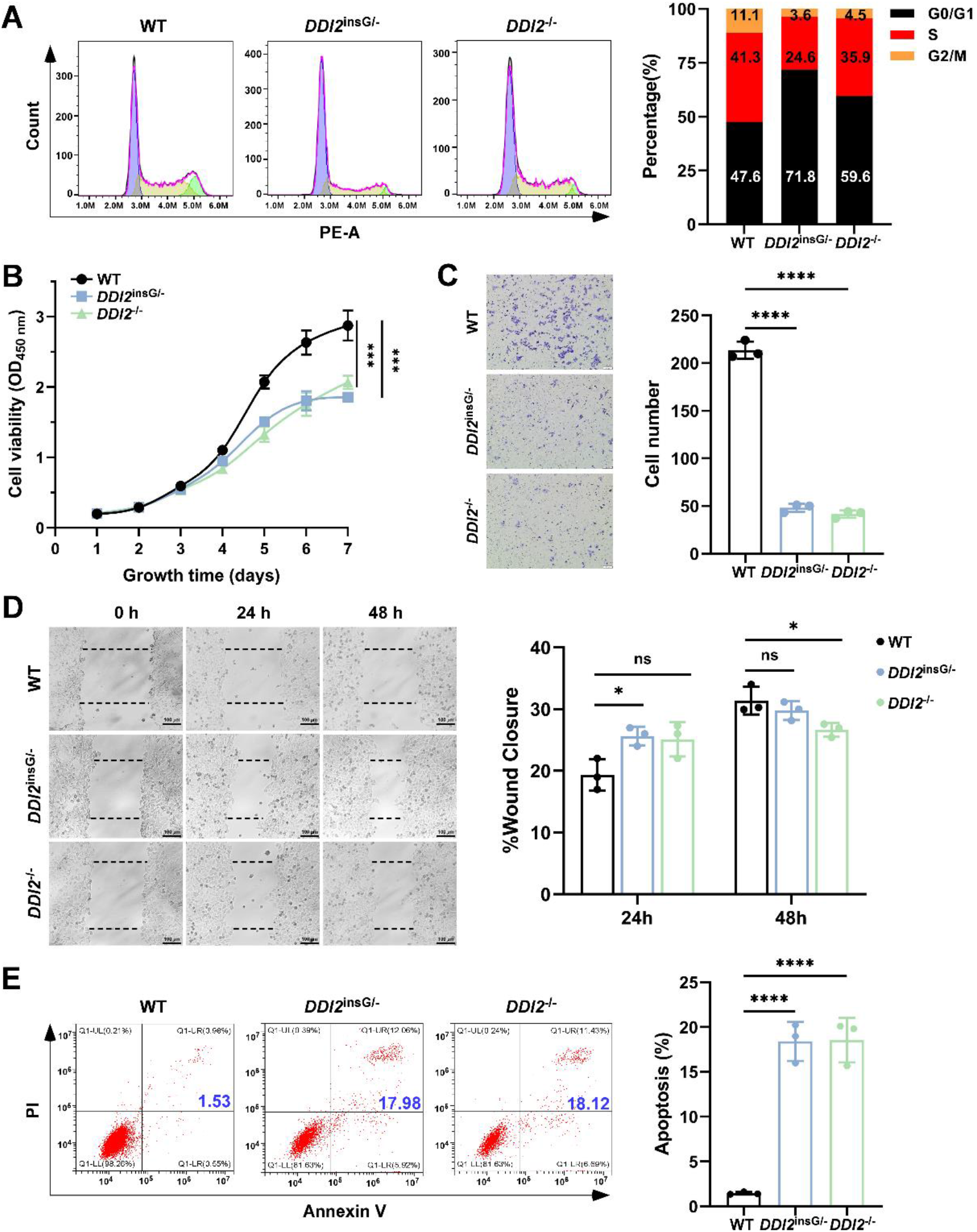
*DDI2* knockout inhibits cell proliferation and migration while promoting cell apoptosis. (A) Flow cytometry was used to detect of cell cycle in wild-type HepG2 cells (WT) and HepG2-derived *DDI2* knockout cells (designated as *DDI2*^insG/-^ and *DDI2*^-/-^). The changes in each stage of the cell cycle were statistically analyzed. (B) A CCK8 assay was conducted to assess the proliferation activity of wild-type HepG2 cells (WT) and *DDI2* knockout cells (*DDI2*^insG/-^ and *DDI2*^-/-^). (C) The transwell assay was employed to evaluate the migration capability of wild-type HepG2 cells (WT) and *DDI2* knockout cells (*DDI2*^insG/-^ and *DDI2*^-/-^), with the number of migrating cells in each group counted. The results are expressed as cell number (mean ± S.D., n = 3), with significant changes indicated (****, p < 0.0001) relative to the WT controls. (D) A cell scratch assay was performed to assess the migration ability of wild-type HepG2 cells (WT) and *DDI2* knockout cells (*DDI2*^insG/-^ and *DDI2*^-/-^), calculating the proportion of scratch healing distance in each group. ns indicates non-significant; *, p < 0.05. (E) Flow cytometry was used to detect cell apoptosis in wild-type HepG2 cells (WT) and *DDI2* knockout cells (*DDI2*^insG/-^ and *DDI2*^-/-^), with the proportion of apoptotic cells statistically analyzed, showing significant differences (n = 3; ****, p < 0.0001).

The results from transwell assay demonstrated that the migratory capacity of liver cancer cells was reduced by *DDI2*^insG/*−*^ or *DDI2*^*−/−*^ (Fig. 3C). Intriguingly, it was found that the inhibitory effects of *DDI2* knockout on wound healing was not significantly pronounced by *DDI2*^*−/−*^ or *DDI2*^insG/*−*^ as deciphered by the cell scratch assay (Fig. 3D). This is much likely to be attributable to the influence of *DDI2* knockout on putative intercellular interactions. As such being the case, further analysis of cell apoptosis unraveled that *DDI2* knockout significantly increased the proportion of apoptotic liver cancer cells (Fig. 3E). To gain sights into such distinct behaviors of *DDI2*-deficient cell*s*, we thus conducted a statistical analysis of the changes in those gene sets related to cell cycle, cell proliferation, and cell death based on the transcriptomic sequencing data. As anticipated, the results unveiled that global alterations in their key genes were consistent with the aforementioned experimental findings (Fig. S4). Altogether, these data indicate that *DDI2* knockout adversely affects the survival of liver cancer cells *in vitro*.

### 3.4. Knockout of *DDI2* inhibits malignant growth of liver cancer cells *in vivo*

Herein, we further examined the *DDI2-*deficient effect of on the *in vivo* growth of liver cancer cells. Initial analysis of those data from the GEEPIA ^22^ and TISIDB ^25^ databases revealed that the expression of DDI2 in liver cancer tissues is higher than that in normal tissues (Fig. 4A). The elevated expression levels of DDI2 were also found to inhibit the abundance of activated CD8^+^ T cells within tumor tissues (Fig. 4B). Collectively, these suggest that high-expressed DDI2 may facilitate the progression of liver cancer *in vivo*. To further investigate this, we conducted subcutaneous tumor-bearing experiments using *DDI2*-knockout liver cancer cells, alongside with their wild-type counterparts, in model nude mice. The observations regarding tumor size, weight, and growth rates indicated that *DDI2* knockout significantly inhibited the growth rate of liver tumors in the nude mice, resulting in notably smaller subcutaneous tumors when compared to those of the control group (Fig. 4, C to E). Subsequently, HE staining (Fig. 4F) and immunohistochemistry analysis (Fig. 4G, *left panels*) of the xenograft tumor tissues revealed increased expression of CASP9 (caspase 9), a key executor enzyme in the apoptosis pathway, in the *DDI2-*deficient tumor group. In contrast, the expression of CCND1 (cyclin D1), a gene associated with cell cycle progression that promotes tumor cell proliferation, was reduced in the *DDI2* knockout tumors (Fig. 4G, *right panels*). Collectively, these findings further support the above notion that *DDI2* knockout can inhibit tumor cell growth *in vivo* by promoting apoptosis and suppressing cell proliferation.

**Figure 4.**
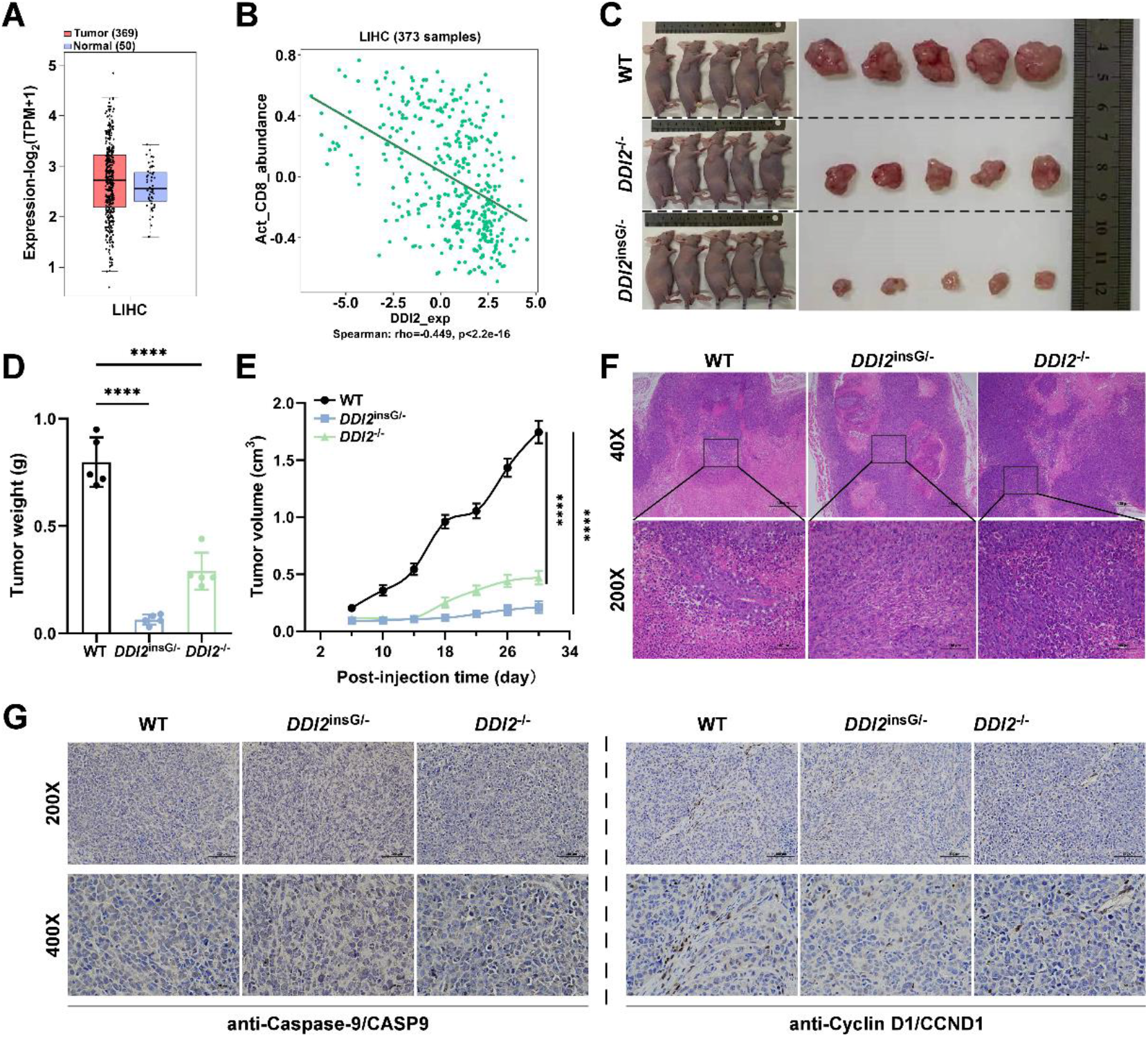
*DDI2* knockout inhibits the malignant growth of cancer cells *in vivo*. (A) A comparative analysis of the differential expression of DDI2 in clinical liver cancer tissues (Tumor) versus adjacent tissues (Normal) was conducted using the GEPIA2 database. (B) The correlation between DDI2 expression and the infiltration level of activated CD8+ T cells in liver cancer tissue was assessed through the TISDIB online database. (C) Tumor formation was observed in wild-type HepG2 cells (WT) and *DDI2* knockout cells (*DDI2*^insG/-^ and *DDI2*^-/-^) following subcutaneous implantation in nude mice. (D) Weight statistics of tumors formed by wild-type HepG2 cells (WT) and *DDI2* knockout cells (*DDI2*^insG/-^ and *DDI2*^-/-^) were recorded from subcutaneous tumor formation experiments in nude mice (n = 5; ***, p < 0.001; ****, p < 0.0001). (E) Growth curves for subcutaneous tumors derived from wild-type HepG2 cells (WT) and *DDI2* knockout cells (*DDI2*^insG/-^ and *DDI2*^-/-^) were plotted. (F) HE staining results of tumors in each group are presented. (G) Immunohistochemical staining analysis the expression of CASP9 and CCND1 in tumor tissues across the different groups.

### 3.5. Restoring DDI2 expression attenuates its deficient inhibitory effect on the survival of liver cancer cells

To further validate the regulatory role of DDI2 in cancer cell survival, we assessed the viability of liver cancer cells following restoration of DDI2 expression in its knockout cells. The results from cell cycle analysis *via* flow cytometry (Fig. 5A) and cell proliferation evaluation using the CCK8 kit (Fig. 5B) demonstrated that the restored expression of DDI2 in *DDI2*^*−/−*^ or *DDI2*^insG/*−*^ cell lines effectively alleviates the suppressive impact of *DDI2* deficiency on hepatoma cell proliferation. Further measurements of intracellular ROS levels indicated that such DDI2 restoration significantly decreases the ROS accumulation caused by *DDI2*^*−/−*^ or *DDI2*^insG/*−*^ (Fig. 5C). These implied that DDI2 enables a marked reduction in intracellular oxidative stress and ensuing damage. Then, assessments of apoptosis levels by flow cytometry revealed that restoring DDI2 expression normalizes the apoptosis levels of its deficient cancer cells to those observed in control cells (Fig. 5D). In addition, we observed that overexpression of DDI2 in wild-type control cells enhanced liver cancer cell proliferation, but concurrently inhibited ROS levels and the apoptosis ratio (Fig. 5, A to D). Taken together, these findings reinforce the notion that *DDI2* knockout indeed suppresses the survival of liver cancer cells, but conversely restoring DDI2 expression effectively counteracts the inhibitory effects of *DDI2* knockout on the growth and survival of these cells.

**Figure 5.**
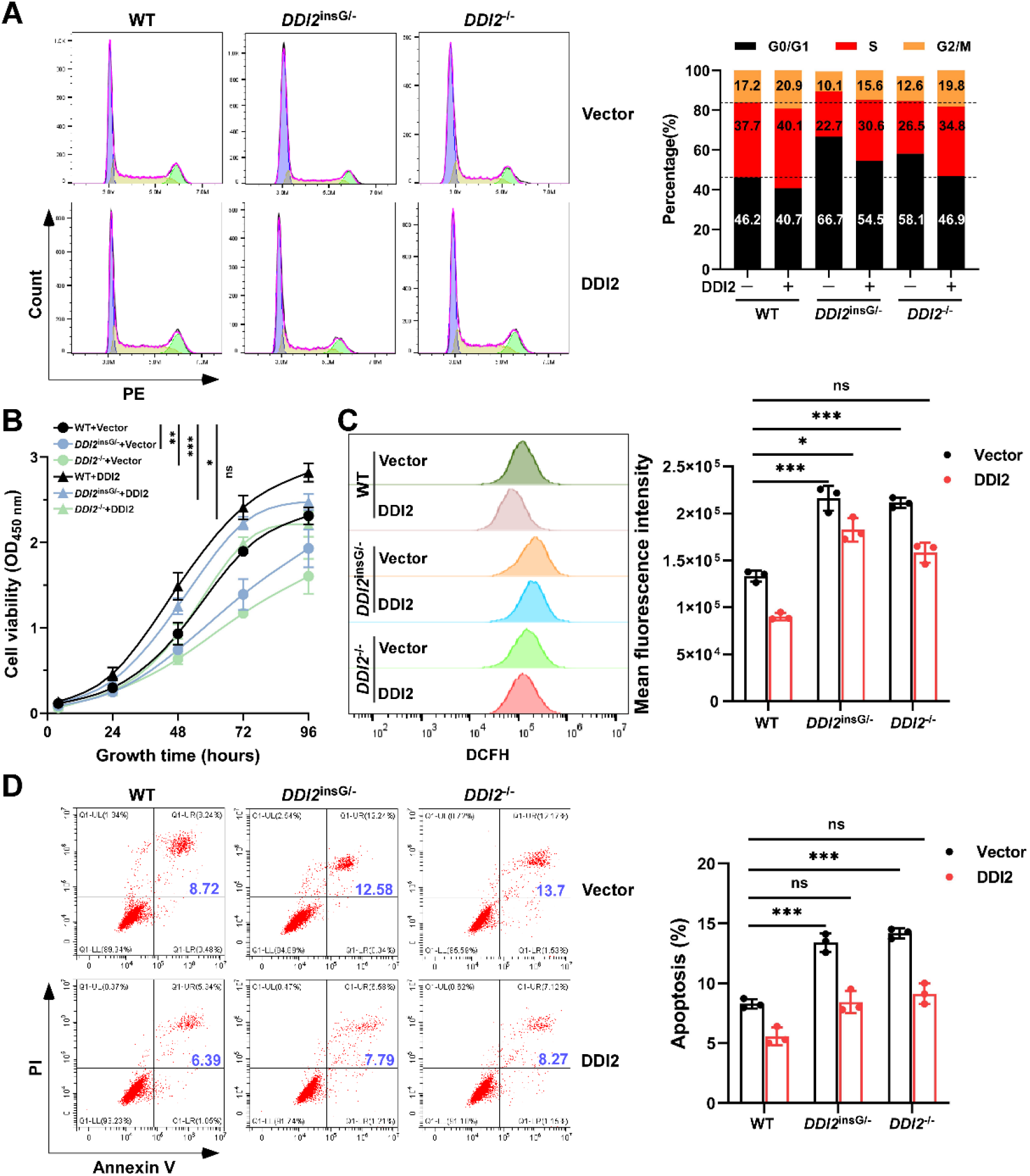
Restoring DDI expression mitigates the inhibitory effect of *DDI2* knockout on cell proliferation and the increased effect of *DDI2* knockout on cell apoptosis. (A) Wild-type HepG2 cells (WT) and HepG2-derived *DDI2* knockout cells (designated as *DDI2*^insG/-^ and *DDI2*^-/-^) were transfected with either a DDI2 expression plasmid or a control vector, followed by an analysis of the cell cycle using flow cytometry. The alterations in each phase of the cell cycle were statistically evaluated. (B) Wild-type HepG2 cells (WT) and *DDI2* knockout cells (*DDI2*^insG/-^ and *DDI2*^-/-^) were similarly transfected and subsequently assessed for proliferation activity using the CCK8 assay. (C) Following transfection with either the DDI2 expression plasmid or the control vector, wild-type HepG2 cells (WT) and *DDI2* knockout cells (*DDI2*^insG/-^ and *DDI2*^-/-^) were analyzed for ROS levels, with the mean fluorescence intensity of ROS calculated (n = 3; ns, non-significant; *, p < 0.05; ***, p < 0.001). (D) Wild-type HepG2 cells (WT) and *DDI2* knockout cells (*DDI2*^insG/-^ and *DDI2*^-/-^) underwent transfection and were evaluated for cell apoptosis through flow cytometry, with the proportion of apoptotic cells statistically analyzed (n = 3; ns, non-significant; ***, p < 0.001).

### 3.6. Restoration of DDI2 rescues from its deficient damages to DNA in liver cancer cells

Here, we further investigated the impact of restoring DDI2 expression on the extents of DNA damage in *DDI2* knockout liver cancer cells. By measuring the fluorescence intensity of the DNA damage marker γH2AX (Fig. 6A) and its protein expression levels (Fig. 6B), it was found that the restoration of DDI2 did not only alleviate the DNA damage caused by *DDI2* knockout to certain levels comparable to those observed in control cells, and also effectively inhibited DNA damage induced by cisplatin. Furtherly, by evaluating changes in the level of another oxidative damage marker, 8-OHdG, we also observed that restoring DDI2 expression significantly reduces the DNA damage exacerbated by *DDI2* knockout (Fig. 6C). In addition, overexpression of DDI2 in wild-type control cells also mitigated putative endogenous DNA damage to certain lower extents in liver cancer cells (Fig. 6, A to C). Altogether, these results demonstrate that DDI2 can negatively regulate the DNA damage process in liver cancer cells, contributing to the survival advantage of liver cancer cells compared to *DDI2* knockout cells.

**Figure 6.**
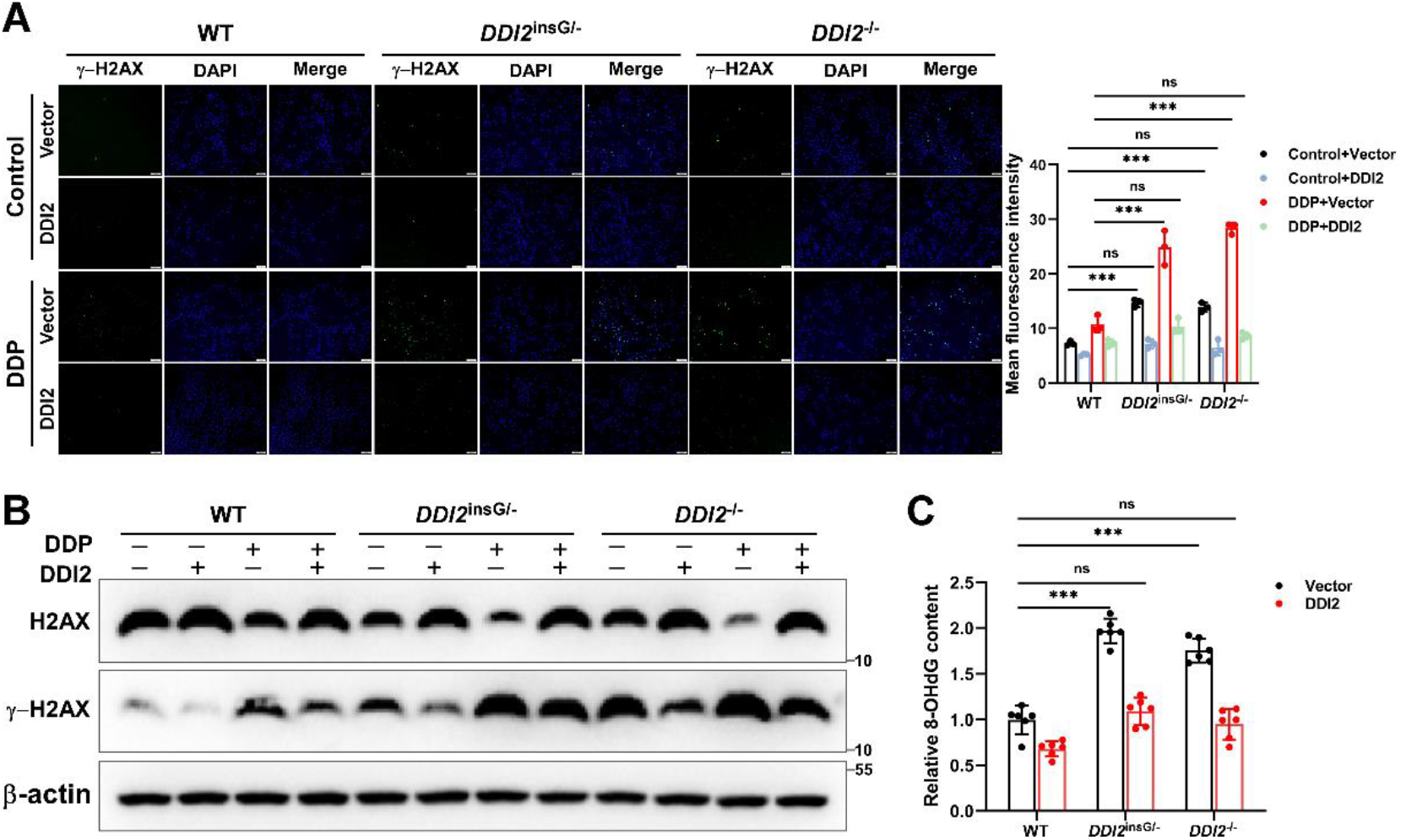
Restoration of DDI2 expression counteracts the promoting effect of *DDI2* knockout on DNA damage. (A) Wild-type HepG2 cells (WT) and HepG2-derived DDI2 knockout cells (designated as *DDI2*^insG/-^ and *DDI2*^-/-^) were transfected with either a DDI2 expression plasmid or a control vector. Following transfection, cells were treated with 10 μM DDP for 22 h and subsequently analyzed for γ-H2AX fluorescence intensity via immunofluorescence. The mean fluorescence intensity of γ-H2AX was calculated and results are expressed as mean fluorescence intensity (n = 3; ns, non-significant; ***, p < 0.001). (B) Wild-type HepG2 cells (WT) and *DDI2* knockout cells (*DDI2*^insG/-^ and *DDI2*^-/-^) underwent transfection with either a DDI2 expression plasmid or a control vector, followed by treatment with 10 μM DDP for 22 h. Western blotting was then conducted to assess the protein levels of H2AX and γ-H2AX. (C) Wild-type HepG2 cells (WT) and *DDI2* knockout cells (*DDI2*^insG/-^ and *DDI2*^-/-^) were transfected with either a DDI2 expression plasmid or a control vector, followed by treated with 10 μM DDP for 22 h, and subsequently analyzed for 8-OHdG content using the 8-OHdG DNA Damage Quantification Direct Kit (n = 6; ns, non-significant; ***, p < 0.001).

## 4. Discussion

The regulation of ROS levels and oxidative DNA damage repair (e.g., by Nrf1 and Nrf2) has emerged as a critical area of research for the development of cancer treatment strategies ^3, 4, 5, 6^. As an upstream protease of Nrf1 processing to become an active antioxidant transcription factor, DDI2 has hence been speculated to play a crucial regulatory role in oxidative stress and ensuing DNA damage, but its specific effects on ROS-leading DNA damage processes in liver cancer cells remain elusive Herein, we have discovered that the knockout of *DDI2* in liver cancer cells inhibits such cell survival and proliferation by elevating ROS levels and exacerbating DNA damage, thereby impeding their malgrowth both *in vitro* and *in vivo* (Fig. 7, *left panel*). Conversely, the restoration of DDI2 alleviates endogenous oxidative stress and DNA damage, effectively counteracting the *DDI2*-deficient inhibitory effects on liver cancer growth (Fig. 7, *right panel*). Collectively, these findings presage that DDI2 is likely to serve as a novel potential target for therapeutic strategies aimed at controlling ROS levels and repairing DNA damage in liver cancer.

**Figure 7.**
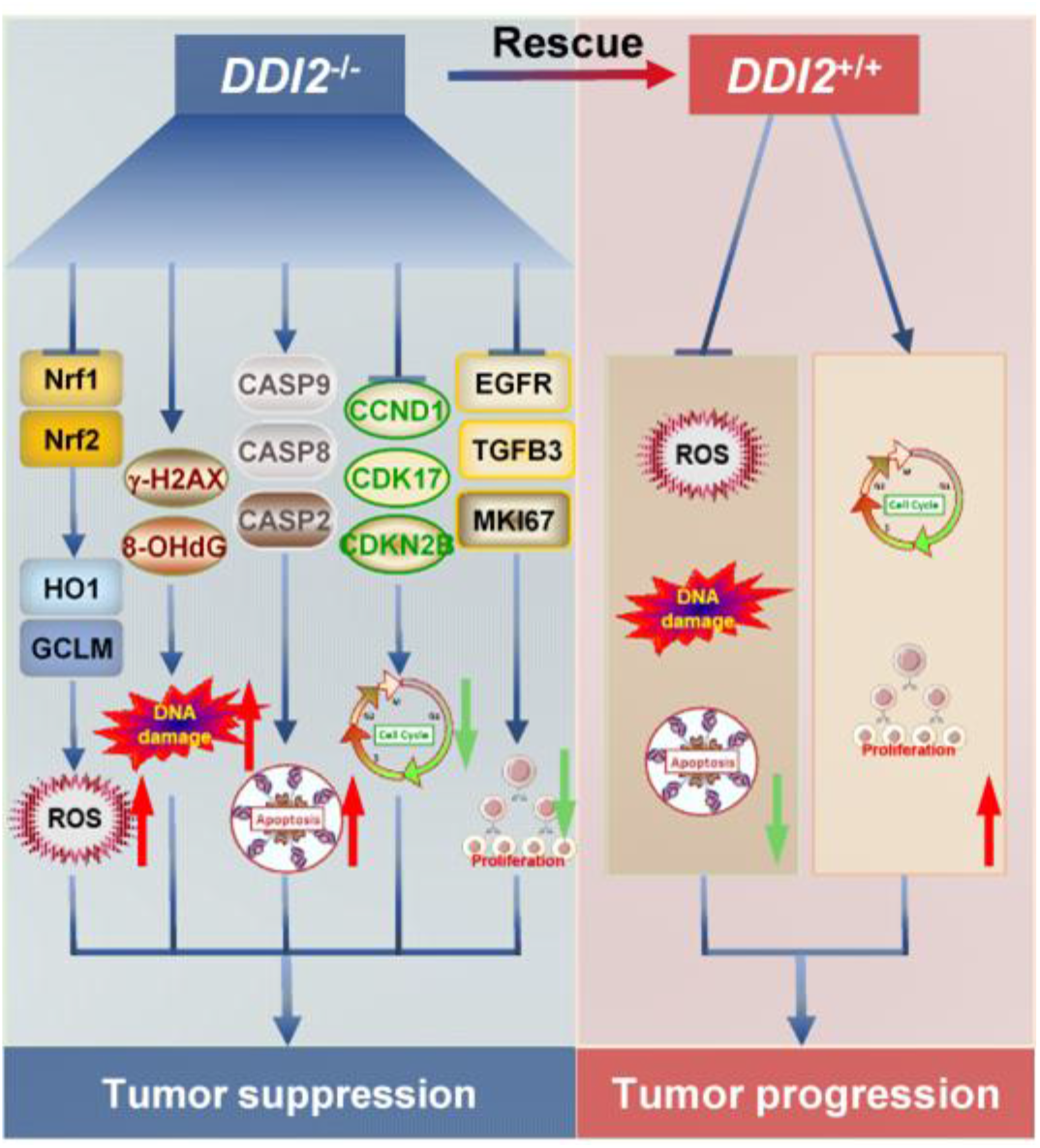
A proposed model for understanding the regulatory effect of DDI2 on liver cancer growth. When DDI2 expression is significantly reduced in liver cancer cells, there is an increase in ROS levels due to the downregulation of antioxidant transcription factors Nrf1 and Nrf2, along with their associated downstream antioxidant genes. This reduction in DDI2 expression also leads to various types of DNA damage and induces cell cycle arrest at the G0/G1 phase. The absence of DDI2 not only suppresses cell proliferation but also promotes apoptosis, ultimately diminishing the in vivo growth of liver cancer cells. Conversely, the restoration or enhancement of DDI2 expression in liver cancer cells inhibits intracellular ROS and DNA damage, thereby promoting cell proliferation and inhibiting apoptosis, which ultimately facilitates the malignant growth of cancer cells.

Currently, DDI2 is recognized to act dually as an aspartic acid protease (to proteolytically process Nrf1) and also as another ubiquitin shuttling factor (to present its cargo to proteasomes regulated by Nrf1) ^13, 26^. Together with its family member DDI1 ^27^, it is also, *de facto*, classified into the ubiquitin receptor protein family. In studies examining the evolutionary development of their structures, DDI2 and DDI1 are frequently referred to as Ddi1-like proteins, which are broadly distributed across animals, plants, fungi, and various protozoan lineages, including apicomplexans, kinetoplastids, and oomycetes. Notably, Ddi1-like proteins have lost their UBA (ubiquitin-associated) domain during their evolutionary transition into vertebrates. In mammalian lineages, the UBA-defective gene underwent duplication, resulting in the emergence of two related proteins containing UBL (ubiquitin-like) and RVP (retroviral protease-like) domains, designated as DDI1 and DDI2 in humans ^26^. The UBL domain is a defining feature of ubiquitin receptor proteins, enabling their binding to ubiquitin (UBQ) and the 26S proteasome, which thus facilitates the recruitment of ubiquitinated substrates for degradation via the proteasome system. The RVP domain is essential for the DDI2’s protease activity, containing the catalytic triad characteristic of aspartic proteases (D[T/S]G) and also playing a role in protein dimerization. Notably, a newly identified helical domain termed HDD, is located adjacent to the RVP in the three-dimensional structure and also exhibits a conserved helical bundle fold ^26, 28^. This structure bears a resemblance to the DNA-binding domain found in transcription factors, and thus enables it for physical binding to DNA damage sites during ubiquitin-dependent proteolysis. This process enables to mediate DDI2’s interaction with potential substrates (e.g., Nrf1) at targeted DNA damage sites, underscoring DDI2’s classification as a novel ubiquitin shuttling factor ^26, 28^, and thus facilitates ensuing repair of damaged DNA responsible for key genes. Therefore, current researches on the function of DDI2 primarily emphasize its capacity to bind ubiquitin substrates and proteasomes, but its role in other biological processes remains to be inadequately understood. Herein, we created two nuanced cell lines of *DDI2* knockout (i.e., *DDI2*^*−/−*^ and *DDI2*^*insG/−*^) from liver cancer, and discovered its biological regulatory functions and potential target genes related to ROS levels, the DNA damage response, and the growth and apoptosis of liver cancer cells (as illustrated in Fig. 7).

As a downstream antioxidant transcription factor of DDI2, Nrf1 has been identified as a key player in maintaining cellular redox balance, as well as cellular glucose, lipid and protein homeostasis, particularly by monitoring the proteolytic activity of proteasome (and this protease *per se*) through mediating the proteasome ‘bounce-back’ response to limited proteasomsal inhibition ^14^. Thereby, DDI2 processes and activates the endoplasmic reticulum membrane-bound Nrf1 to because an active CNC-bZIP factor during the Nrf1-mediated proteasome ‘bounce-back’ reaction, facilitating the entry of the active isoform of Nrf1 into the nucleus to regulate proteasomal transcriptional expression and its *de novo* activity. Upon the knockout of *DDI2*, examined cells exhibit heightened sensitivity to proteasome inhibitors, resulting in the accumulation of high molecular weight, ubiquitinated proteins (e.g., of Nrf1) by putative inefficiency of the ubiquitin-proteasome system (UPS) ^13, 14^. Thus, DDI2 indeed plays a crucial role in regulating both cellular redox stability and even ROS levels. This is further supported by our discovery, for the first time, that the knockout of *DDI2* can downregulate Nrf1 and Nrf2 at both mRNA and protein levels, thereby leading to an imbalance in redox status with increased ROS levels in liver cancer cells (Figs. 1 and S2).

In such oxidative and adverse conditions with induced DNA damage to high levels, the DNA replication process is disrupted, leading to slowed DNA synthesis and/or stalled replication forks, a phenomenon known as DNA replication stress. This stress is accepted as a primary contributor to genomic instability and also as a hallmark of both precancerous and cancerous cells ^29, 30.^ To address this issue, all cellular life forms have evolved multiple signaling pathways and defense processes to counteract DNA damage and replication stress ^31^. During such DNA replication, replication termination factor 2 (RTF2) is present at stalled replication forks, whilst DDI2 can facilitate the induction of RTF2 to release from stalled replication forks, and thus its removal can alleviate replication stress, thereby maintaining genomic stability ^18^. However, the absence of DDI2 can lead to defects in the replication fork restart, hyper-activation of DNA damage signals, accumulation of single-stranded DNA (ssDNA), increased sensitivity to replication drugs, and chromosomal instability ^18^. Some studies have indicated that microRNA-3607 can target DDI2 to regulate the DNA damage repair pathways, potentially interfering with the malignant behavior of colorectal cancer cells ^17^; however, such a regulatory effect of DDI2 on DNA damage had not been thoroughly explored in this context. Notably, another research had further shown that DDI2 can function as a key repair enzyme of DNA–protein crosslink (DPC, a common form of DNA damage that disrupts DNA replication, repair, transcription, and recombination) ^32^. Collectively, these findings suggest that DDI2 plays a significant role in regulating the DNA damage repair processes. This is evidenced by a recent report indicating that DDI2 can promote the metastasis of colon cancer ^33^, but its regulatory effects and mechanisms remain unclear. In the present study, we have provided the evidence demonstrating that DDI2 indeed enhances the survival of liver cancer cells by promoting DNA damage repair.

In conclusion, here we have, for the first time, discovered that DDI2 acts dually as an important regulator of both oxidative stress defense and DNA damage repair, such that it can promote the growth and survival of liver cancer cells both *in vitro* and *in vivo* by monitoring intracellular ROS levels and inhibiting the aggravation of DNA damage. Overall, this study provides further insights into the biological function of DDI2 insofar as to enable it to be developed as a novel potential strategic target for preventing liver cancer.

## Author contributions

K.L., S.H., W.R., Q.Z. and L.Q. performed all the experiments with help of M.W., collected all the relevant data, and made the manuscript draft with figures and supplemental information. M.W. did bioinformatics analysis of datasets. As a native English speaker, Z.Z. helped to polish the English language and edited this manuscript. Lastly, Y.Z. together with M.W. designed and supervised this study, analyzed all the data, helped to prepare all figures with cartoons, wrote and revised the paper.

## Acknowledgments

We thank to all those present and past members of Prof. Zhang’s laboratory (at Chongqing University, China) for giving critical discussion and invaluable help with this work.

## Data Availability Statement

All data needed to evaluate the conclusions in the paper are present in this publication along with the supplementary documents that can be found online. Additional other data related to this paper may also be requested from the corresponding author (with a lead contact at the, or).

