## Supplemental FigureS1-S4 and Table S1 for "Deficiency of DDI2 suppresses liver cancer progression by worsening cell survival conditions"

#### Figure S1

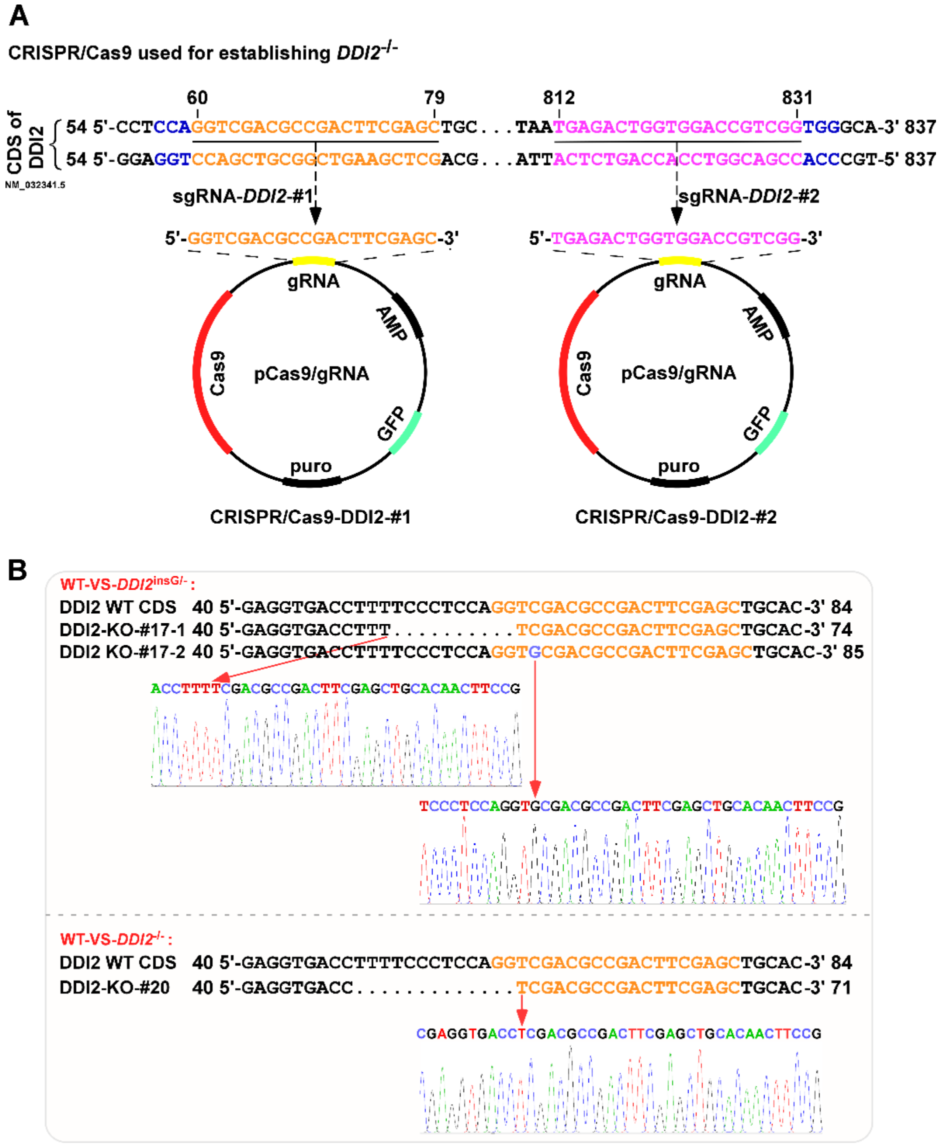


**Figure S1. Establishment and sequence validation of *DDI2* knockout cell lines.** (A) Construction of two CRISPR/Cas9-DDI2 plasmids. (B) Sequence alignment of the *DDI2* gene knockout sites in two *DDI2* knockout monoclonal cell lines and the wild-type (WT) cell line, along with the sequencing peak map of the knockout sites in DDI2-KO-#17 (*DDI2*$^{isnG/-}$) and DDI2-KO-#20 (*DDI2*$^{-/-}$) monoclonal cells.

#### Figure S2

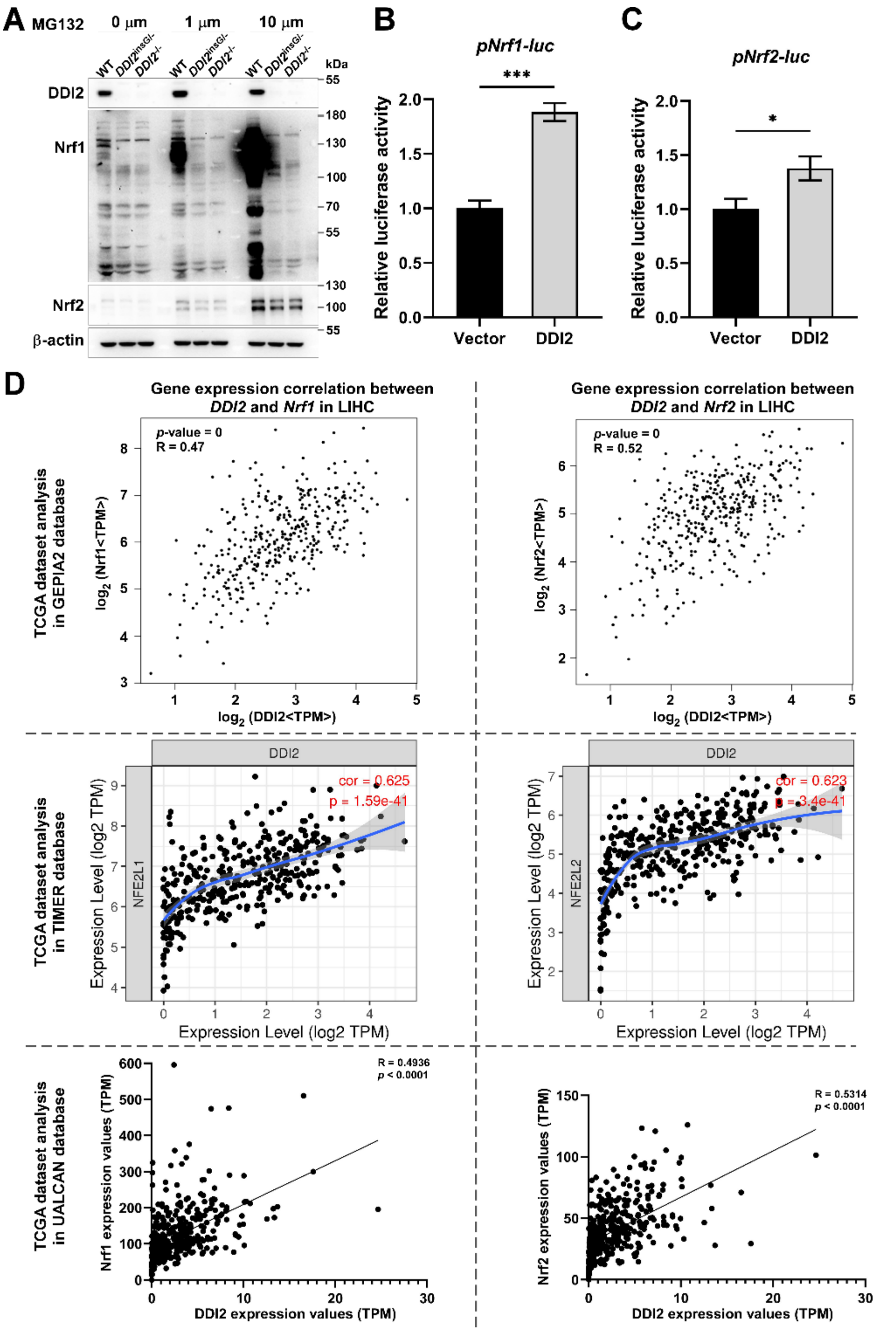


**Figure S2. The regulatory effects of DDI2 on Nrf1 and Nrf2, as well as their expression correlation in liver tissue.** (A) Wild-type HepG2 cells (WT) and *DDI2* knockout cells (*DDI2*$^{isnG/-}$ and *DDI2*$^{-/-}$) were treated with 0, 1 or 10 μM MG132 for 16 h. Subsequently, Western blotting was performed to evaluate the protein levels of DDI2, Nrf1 and Nrf2. (B, C) HEK293T cells were transfected with either the promoter pNrf1-luc (B) or pNrf2-luc (C), along with the DDI2 expression plasmid, and then subjected to a dual luciferase reporter gene detection assay (*, $p < 0.05$; ***, $p < 0.001$). (D) The correlation of gene expression between DDI2 and Nrf1 or Nrf2 was analyzed using the TCGA-LIHC dataset from the GEPIA2, TIMER and UALCAN databases.

### Figure S3

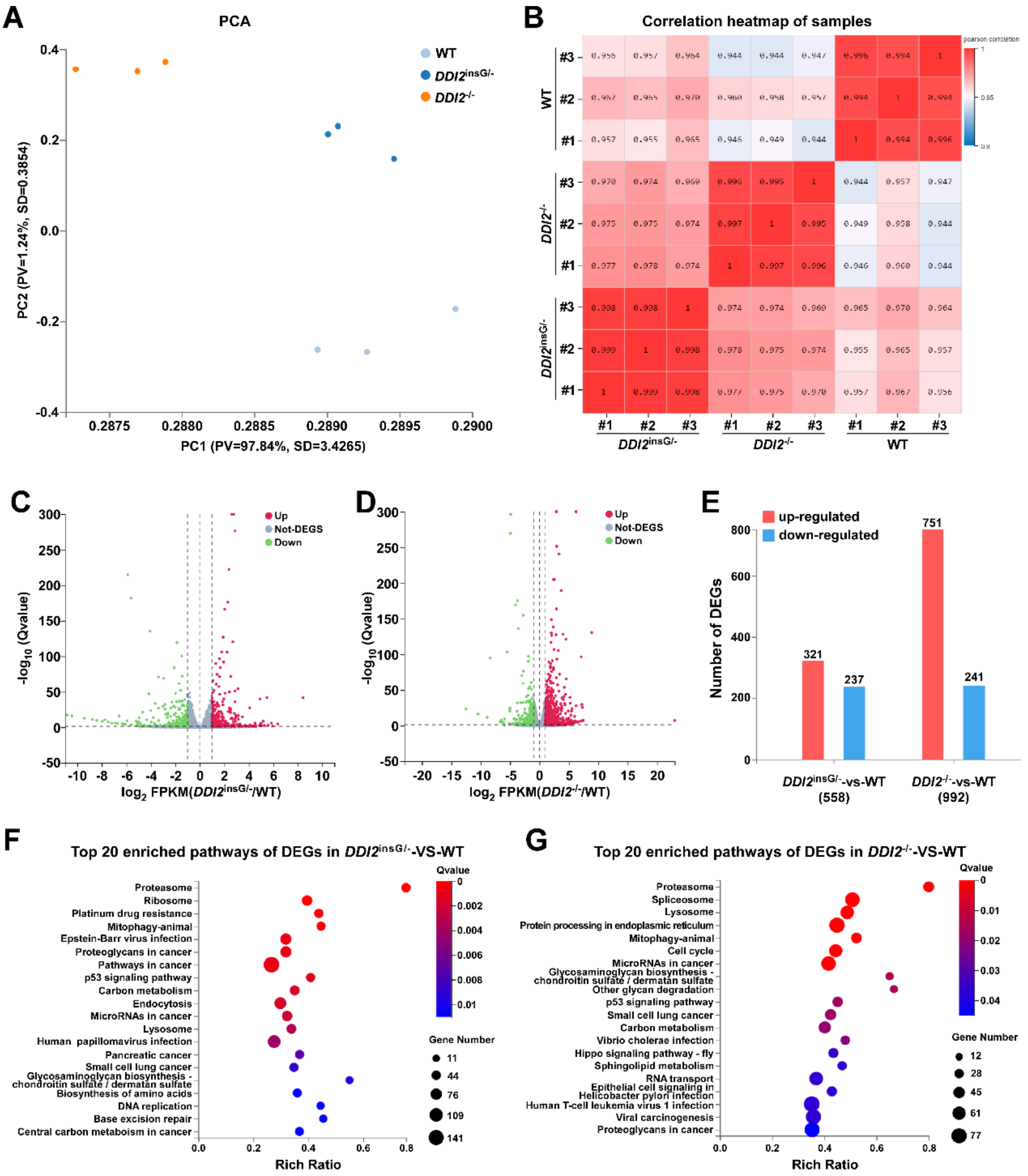


**Figure S3. Global analysis of transcriptome sequencing results.** (A, B) Wild-type HepG2 cells (WT) and *DDI2* knockout cells (*DDI2*$^{isnG/-}$ and *DDI2*$^{-/-}$) underwent transcriptome sequencing, with each group comprising three biological replicates. Subsequently, principal component analysis (PCA) (A) and correlation of sequenced samples (B) were performed. (C, D) Following statistical analysis of differentially expressed genes resulting from knockout of *DDI2* in HepG2 cells, volcano plots illustrating gene expression trends in *DDI2*$^{isnG/-}$ cells(C) and *DDI2*$^{-/-}$ cells (D) were generated, comparing them to the gene expression levels in WT cells. (E) A histogram depicting the number of differentially expressed genes in each group is presented. (F, G) The top 20 pathways enriched with differentially expressed genes in *DDI2*$^{isnG/-}$ (F) and *DDI2*$^{-/-}$ (G) cells were analyzed in comparison to WT cells.

#### Figure S4

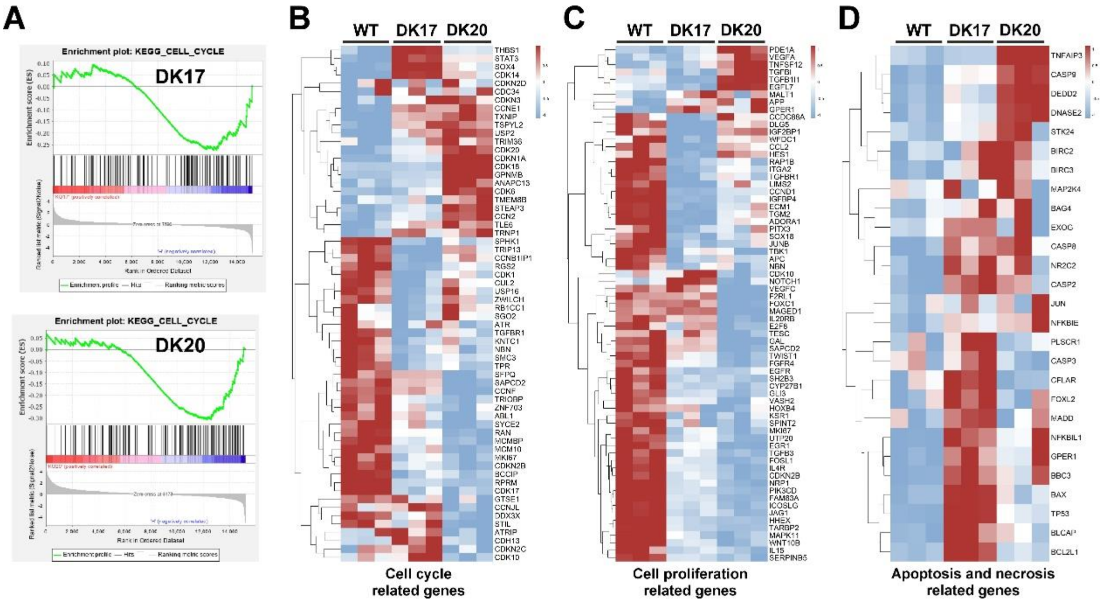


**Figure S4. Changes in cell cycle, proliferation, and apoptosis-related genes following *DDI2* knockout.** (A) Gene Set Enrichment Analysis (GSEA) of the cell cycle pathway in $DDI2^{isnG/-}$ (NES = -1.275, p = 0.019) and $DDI2^{-/-}$ (NES = -1.392, p = 0.0011) cells was conducted in comparison to wild-type HepG2 cells (WT), utilizing transcriptome sequencing data. (B-D) Differential expression heatmaps of genes related to cell cycle, cell proliferation, apoptosis and necrosis were generated for wild-type HepG2 (WT) cells and DDI2 knockout cells ($DDI2^{isnG/-}$ and $DDI2^{-/-}$) based on the transcriptome sequencing analysis.

**Table S1. The key reagents and resources used in this study**

| Reagents or resources | Identifier | Source |
|---|---|---|
| **1. Oligonucleotides for qPCR** | | |
| β -actin FW | CATGTACGTTGCTATCCAGGC | Tsingke |
| β -actin REV | CTCCTTAATGTCACGCACGAT | Tsingke |
| Nrf1 FW | TGGAACAGCAGTGGCAAGATCTCA | Tsingke |
| Nrf1 REV | GGCACTGTACAGGATTTCACTTGC | Tsingke |
| Nrf2 FW | TCAGCGACGGAAAGAGTATGA | Tsingke |
| Nrf2 REV | CCACTGGTTTCTGACTGGATGT | Tsingke |
| HO1 FW | TCAAAAAGATTGCCCAGAAAGCC | Tsingke |
| HO1 REV | GGTAGAGCTGCTTGAACTTGGTG | Tsingke |
| GCLM FW | GTGTGATGCCACCAGATTTGAC | Tsingke |
| GCLM REV | CACAATGACCGAATACCGCAGT | Tsingke |
| DDI-2 FW | TCCAGTGCAGTTCCCAAACTTAC | Tsingke |
| DDI-2 REV | ATGCTGCTCACCGTGTACTGTGTGC | Tsingke |
| PSMB5 FW | TCGGCAATGTCGAATCTATGAGC | Tsingke |
| PSMB5 REV | ATGGCTGGGGGCGCAGCGGATTGCA | Tsingke |
| PSMB6 FW | TCAAGAAGGAGGGCAGGTGT | Tsingke |
| PSMB6 REV | AGACTTCTACAACGATCCCCTCT | Tsingke |
| PSMB7 FW | GATACAAGAGCAACTGAAGGGATG | Tsingke |
| PSMB7 REV | ATGGCGGCTGTGTCGGTGTATGCTC | Tsingke |
| **2. Antibodies** | | |
| Nrf1 | Made by us | Zhang's Lab |
| Nrf2 | ab62352 | Abcam |
| β-actin | TA-09 | ZSGB-BIO |
| HO1 /HMOX1 | ab68477 | Abcam |
| GCLM | ab126704 | Abcam |
| H2AX | ab124781 | Abcam |
| γ -H2AX | ab81299 | Abcam |
| DDI-1 | H00414301-B01P | Novus Biologicals |
| DDI-2 | A304-630A-T | Bethyl |
| PSMB5 | ab3330 | Abcam |
| PSMB6 | ab150392 | Abcam |
| PSMB7 | ab154745 | Abcam |
| **3. Chemicals** | | |
| DDP | 15663-27-1 | Aladdin |
| MG132 | M7449 | Sigma Aldrich |
| NAC | A601127 | Sangon Biotech |
| DMSO | 67-68-5 | Aladdin |
| **4. Oligonucleotides for expression constructs** | | |
| Nrf1-LUC-#1 FW | CCTAGGCCTGCTAGCGCGACTGAG TTTGTCTCTACACCT | Tsingke |

| | | |
|---|---|---|
| Nrf1-LUC-#1 REV | CTTCAGAGAAAAGCTTGCTGAAGG ACCAGAATGTTTATGCT | Tsingke |
| Nrf2-LUC FW | CCAGGAGTTTGGTACCAGCCTGGG CAACATAGTGA | Tsingke |
| Nrf2-LUC REV | CCAGCTCCAAGTAGATCTTGATGA GCTGTGGA | Tsingke |
| DDI2 FW | CTAGCTAGCATGCTGCTCACCGTG TACTGTGTGCGGAGG | Sangon Biotech |
| DDI2 REV | CGGAATTCTCATGGCTTCTGACGC TCTGCATCCTCTGC | Sangon Biotech |
| CRISP/Cas9-DDI2-1-F | AAACACCGGCTCGAAGTCGGCGTC GACC | Tsingke |
| CRISP/Cas9-DDI2-1-R | CTCTAAAACGGTCGACGCCGACTT CGAGC | Tsingke |
| CRISP/Cas9-DDI2-2-F | AAACACCG TGAGACTGGTGGACCGTCGG | Tsingke |
| CRISP/Cas9-DDI2-2-R | CTCTAAAAC CCGACGGTCCACCAGTCTCA | Tsingke |
| **5. Recombinant DNAs** | | |
| pcDNA3.1 | V79020 | Invitrogen |
| pGL3-Basic | VQP0121 | Promega |
| pRL-TK | VQP0126 | Promega |
| **6. Software and Algorithms** | | |
| Canvas X | https://www.canvasgfx.com/ | Canvas GFX, Inc. |
| Chromas 2.4.1 | http://technelysium.com.au/wp/chromas/ | Technelysium Pty Ltd. |
| Excel | https://www.microsoft.com/ | Microsoft |
| Primer Premier 5 | https://www.PremierBiosoft.com/ | PREMIER Biosoft International |
| CFX Manager 3.1 | https://bio-rad-cfx-manager.com/ | Bio-Rad |
| R | version 4.3.0 | |
| GSEA | version 4.0.3 | |
| **7. Others** | | |
| Cas9/gRNA Construct Kit | VK001 | v-solid |
| Dual-luciferase reporter assay system | E1910 | Promega |
| RNAsimple Total RNA Kit | DP419 | Tiangen Biotech |
| Lipofectamine®3000 Transfection Kit | L3000-015 | Invitrogen |
| GoTaq®qPCR Master Mix | A6001 | Promega |
